# CandyCollect Open to Closed (O2C) Microfluidic System for Rapid and User-Centric Detection of Group A Streptococcus

**DOI:** 10.1101/2025.02.10.635313

**Authors:** Kelsey M. Leong, J. Carlos Sanchez, Cosette Craig, John A. Tatka, René R. Arvizu, Ingrid H. Robertson, Victoria A. M. Shinkawa, Timothy R. Robinson, Megan M. Chang, Xiaojing Su, Sanitta Thongpang, Ashleigh B. Theberge, Erwin Berthier, Ayokunle O. Olanrewaju

## Abstract

We created CandyCollect, a lollipop-inspired, child-friendly saliva sampling device that provides enhanced comfort compared to common throat and mouth swabs. The device has an open microchannel and a functionalized surface that effectively captures and stores the saliva and pathogens. We recently demonstrated the ability to combine the CandyCollect with rapid antigen detection tests (RADTs) for fast and accessible detection of group A streptococcus (GAS).

However, the current procedure necessitates manual steps to elute and transfer captured antigens from the lollipop to RADTs, which relies on the user to follow complex instructions, increasing the likelihood of error. In this work, we developed the CandyCollect Open to Closed (O2C) microfluidic system to automate antigen elution and streamline detection with RADTs. The O2C system securely seals the open microchannels of the CandyCollect device to create a continuous closed channel for seamless delivery of elution reagents to release antigens at the press of a button for subsequent lateral flow detection. Our system leverages and combines the unique benefits of both open channel and closed channel microfluidics into one platform, further enhancing the potential of both methods. We performed experiments with GAS bacteria spiked into saliva and compared the manual and O2C elution methods. Both the manual and O2C methods showed clear test lines on the RADT strips at clinically relevant GAS concentrations (between 5 ×10^5^ and 10^9^ CFU/mL). The O2C provided comparable or better results than the manual procedure in a more convenient form factor. The O2C platform has the potential to enable user-friendly screening for respiratory pathogens by minimally trained users in decentralized settings.

## Introduction

Oral samples for the diagnosis of respiratory diseases are critical diagnostic tools for children as certain bacterial and viral diseases can lead to long-term debilitating conditions or poor prognosis when left untreated.^1–3^ However, the lack of patient-centric access to reliable, easy-to-use diagnostic testing is a barrier that limits early and patient-driven testing. Furthermore, pediatric samples are particularly challenging to obtain as children are frequently uncooperative with procedures that cause pain, trauma, or anxiety, and parents sometimes put off procedures that induce pain and discomfort to their child, which can hinder early detection and intervention measures. Methods such as the pharyngeal swab, sputum collection, and nasal swab are unpleasant for both adults and children,^4^ and pharyngeal swabs and sputum collection are also challenging to perform in a home setting.^5^ Child-friendly diagnostics are critical for diseases like streptococcal pharyngitis, commonly known as ‘strep throat’, influenza, respiratory syncytial virus (RSV), and pneumonia that affect millions of children worldwide, occasionally resulting in severe and long-term consequences.^1^

We previously developed a lollipop-inspired device termed ‘CandyCollect’ that enables user-friendly, particularly child-friendly, high-quality pathogen collection from saliva using open microfluidic channels and functionalized surfaces.^6–8^ Preliminary results show that CandyCollect captures a variety of microbes and is preferred to traditional throat swabs, oral swabs, and spit tubes.^6^ In a clinical study of children with group A streptococcus (GAS) pharyngitis, samples were collected with both CandyCollect and traditional mouth swabs. After analysis with quantitative polymerase chain reaction (qPCR), all participants (30/30) were positive for GAS for both sampling methods demonstrating that CandyCollect can be a feasible oral swab alternative.^8^ Furthermore, we demonstrated that CandyCollect can be used in place of a traditional pharyngeal swab with a commercial GAS rapid antigen detection test (RADT) kit, expanding the testing scenarios where this device can be applied.^9^

While our initial results integrating CandyCollect with RADTs were promising, the current workflow requires manual elution steps to transfer pathogens from the open microchannels on the CandyCollect lollipop to the RADTs. These manual handling steps rely on the user to follow complex instructions, increasing the likelihood of error during testing and limiting the user-friendliness of this platform. To enable at-home use of CandyCollect, there is a need for a more streamlined device that minimizes user interaction and automates key extraction and testing steps.

Currently, there are a few sampling devices available that are similar to our device but lack some of the features that make the CandyCollect unique. Self-LolliSponge™, V-check COVID-19, and Whistling COVID-19 are all saliva collection devices that facilitate self-collection of saliva with a minimally invasive protocol,^10,11^ however these devices lack the patient-centric candy aspect of the CandyCollect and there is a need for further integration of these devices with RADTs to enable point-of-care testing.

We recently developed the prototype CandyCollect Open-to-Closed (O2C) microfluidic system which aims to integrate pathogen capture, inline elution, and RADT detection into a single platform. The concept of integrating closed and open microfluidic channels has been explored before^12,13^ where continuous flow is provided by the closed channels and an intermediate channel opening provides access to fluids for sampling or measurement. Our approach takes a different route by leveraging the open microchannels to capture saliva and pathogens, while the closed microchannels deliver and transport reagents. By integrating these complementary functions into a single system, we maximize the unique capabilities of both microfluidic designs. The CandyCollect O2C device includes a top layer with a closed microfluidic channel, an intermediate device ‘ceiling’ layer to prevent leakage, and a bottom layer with a through-hole for liquid output. The bottom and top layers enclose the CandyCollect effectively creating a tight ‘sandwich’ that converts the open pathogen collection channel into a closed channel for inline testing. Using a finger pump,^14^ pathogen elution reagents are flushed through the now closed CandyCollect, and out the bottom where the sample can be collected and delivered to a RADT for testing and readout. Our results show that the O2C provides similar or better performance to the manual sample elution method, while also streamlining elution steps, decreasing exposure to potential infectious pathogens, and enabling simple manipulation of smaller sample volumes. The system combines a user-friendly sampling method with a simplified testing process which lays the foundation for future developments of this prototype into a robust patient-centric diagnostic platform for decentralized settings.

## Results and Discussion

### Components and assembly of CandyCollect O2C device

The CandyCollect was recently validated as a more comfortable alternative to the manufacturer-provided pharyngeal swab in the Areta Strep A Swab Test™.^9^ We found that when the CandyCollect was substituted into the workflow of the RADT kit, positive test results were detected by eye and with a signal image quantification program for bacterial concentrations as low as the manufacturer’s “minimal detection limit” - 1.5 × 10^5^ CFU/mL;^9^ demonstrating that the CandyCollect can be used with LFA rapid tests.

Here we engineered the O2C system to automate multiple user steps required in the manual RADT elution procedure. The open channels of CandyCollect enable capture and storage of pathogens but discourage controlled fluid delivery and automated manipulation of the sample. We transformed the open channels of CandyCollect to closed channels by placing a device ‘ceiling’ layer over the channels. A small hole in the ceiling layer at the inlet of the CandyCollect allowed reagents from the top channel to flow through the spiral of the CandyCollect. When the O2C device encloses the CandyCollect stick, the pressure of the top layer pushes down on the ceiling layer effectively and securely enclosing the open microchannels. Once the system is assembled, pressing the finger pump facilitates user-friendly addition of elution reagents to the CandyCollect to release captured antigens. The O2C device **(Figure 1A)** consists of two 3D-printed layers **(Figure 1A(ii)&(vi))**, a silicone finger pump retrofitted from an e-cigarette cap^14^ **(Figure 1A(i))**, and a silicone ceiling layer **(Figure 1A(iv))** situated on top of the CandyCollect **(Figure 1A(v))**. The top layer contains a 3D-printed microfluidic channel sealed with tape to retain elution reagents; upstream of the reagent channel is an inset for the finger pump attachment. The bottom layer has an inset for the CandyCollect device with a through-hole in the center for liquid elution into a reservoir. When all components are assembled and aligned, the outlet of the reagent channel aligns with the inlet of the CandyCollect microchannel **(Figure 1B)**, creating a closed, continuous pathway for liquid to travel through.

**Figure 1:**
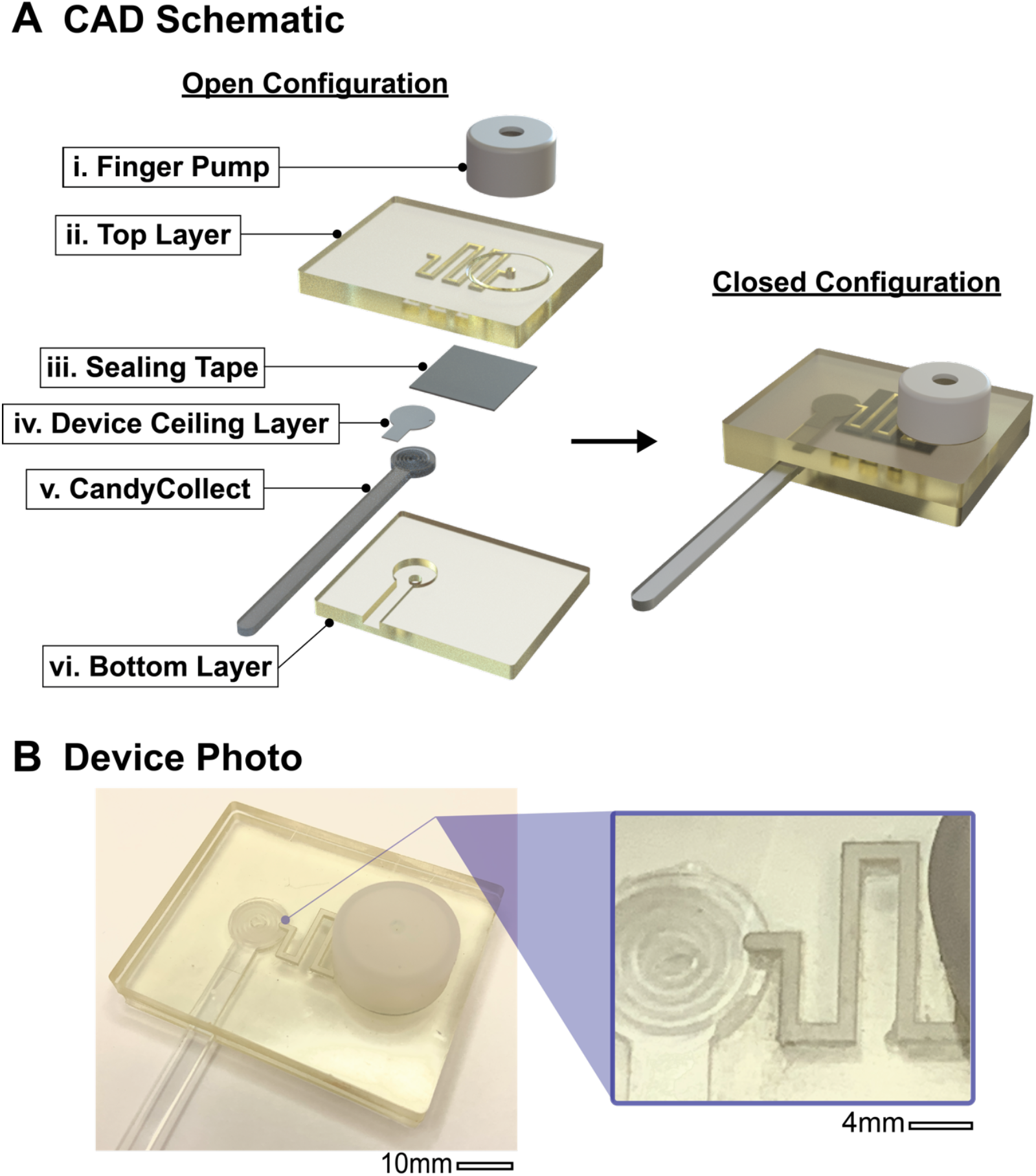
Design and assembly of the O2C system for automated elution of antigens. **A)** Exploded CAD renderings of the O2C components. The rendering on the left contains a CandyCollect which has open channels and has yet to be sealed. The rendering on the right shows the entire system when it is assembled and the microchannels on the CandyCollect are fully enclosed. **B)** Image of the 3D-printed O2C device with fully assembled components. The inset to the right highlights the connection between the reagent channel on the top layer of the O2C and the CandyCollect.

### Simplified antigen elution workflow using CandyCollect O2C

The O2C automates the mixing of elution reagents with potentially infectious agents, enabling controlled and direct interaction of reagents with the captured pathogens in the microchannel. Mixing and elution into a reservoir occur consecutively with the single press of the finger pump. To mimic the sampling process, we first inoculated the CandyCollect with saliva and bacteria for 10 minutes **(Figure 2A(i), red food coloring used for visualization)**. Elution reagents are pipetted into the top channel **(Figure 2A(ii), blue food coloring used for visualization)**. The device components are assembled and held tightly together by binder clips to prevent leakage **(Figure 2A(iii))**. Our current iteration uses binder clips for frugal prototyping; however, these can be exchanged for other clamping mechanisms in future designs to facilitate usability and manufacturability.^15,16^ The finger pump is pressed (**Figure 2A(iv))**, and the liquid elutes through the CandyCollect channel and out into a reservoir **(Figure 2A(v)). Figure 2B** shows side and top views of the liquid path through the device. Elution buffer **(blue)** is flushed through the reagent channel, into the CandyCollect channel **(red)**, mixing together, before being ejected into the reservoir **(dark purple droplet)** where the LFA test is performed. The current design enables automated mixing and elution; in future design iterations this workflow can be improved by including a holder for the lateral flow strip which can be directly attached to the O2C for minimal contact with potentially infectious material.

**Figure 2:**
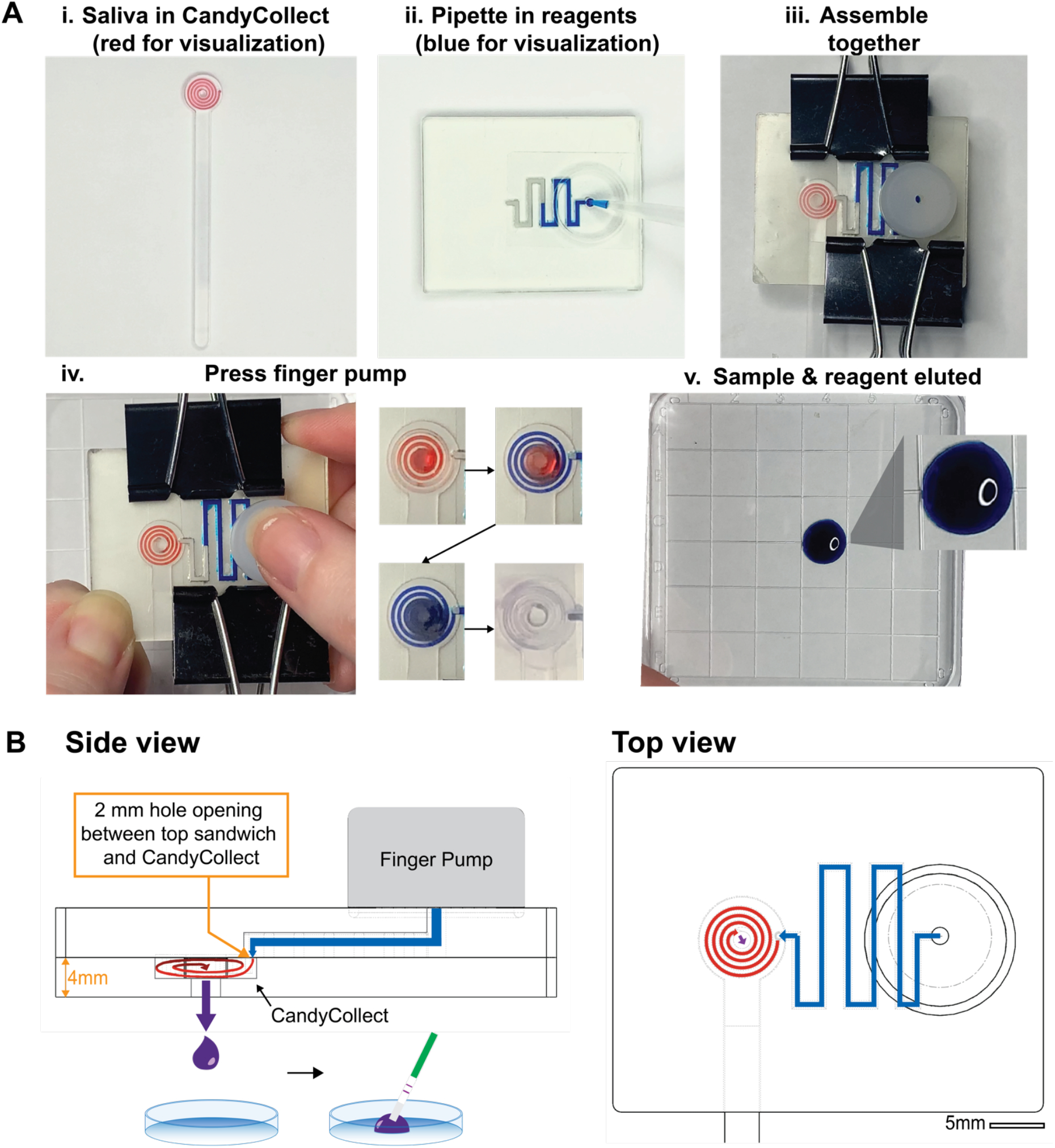
Proof-of-concept operation of the CandyCollect O2C system using food dye solutions. **A)** Workflow for proof-of-concept testing with the CandyCollect O2C device. i) Inoculate CandyCollect with bacteria for 10 minutes (red food coloring for visualization). ii) Pipette elution buffer into reagent channel in the top layer (blue food coloring for visualization). iii) Stack components and clamp together to ensure tight seal between all layers. iv) Attach finger pump and press to elute. The time series photos depict the progression of liquid elution as the pump is pressed down. v) The sample is eluted from the CandyCollect open fluidic channel into reservoir. The inset shows a zoomed in photo of the intact droplet. **B)** Side view and top view of the device with fluid pathway highlighted in blue *(elution buffer)* and red *(saliva with bacteria)*. When the finger pump is pressed, the two liquids mix together *(depicted in dark purple)* and output through the bottom layer outlet into a petri dish for analysis with a lateral flow strip.

### Testing the CandyCollect O2C with clinically relevant concentrations of GAS in spiked saliva

For our initial proof-of-concept experiments, a previously optimized total elution reagent volume of 200 µL^9^ was used for both the manual and microfluidic elution methods. GAS concentrations between 5 ×10^5^ and 10^9^ CFU/mL showed clear test lines on the LFA strip (**Figure 3A and S1**), and the quantified test line signals were above the positivity threshold (**Figure 3C**). As concentrations of GAS increased, the O2C method generally provided higher signal than the manual method.

**Figure 3:**
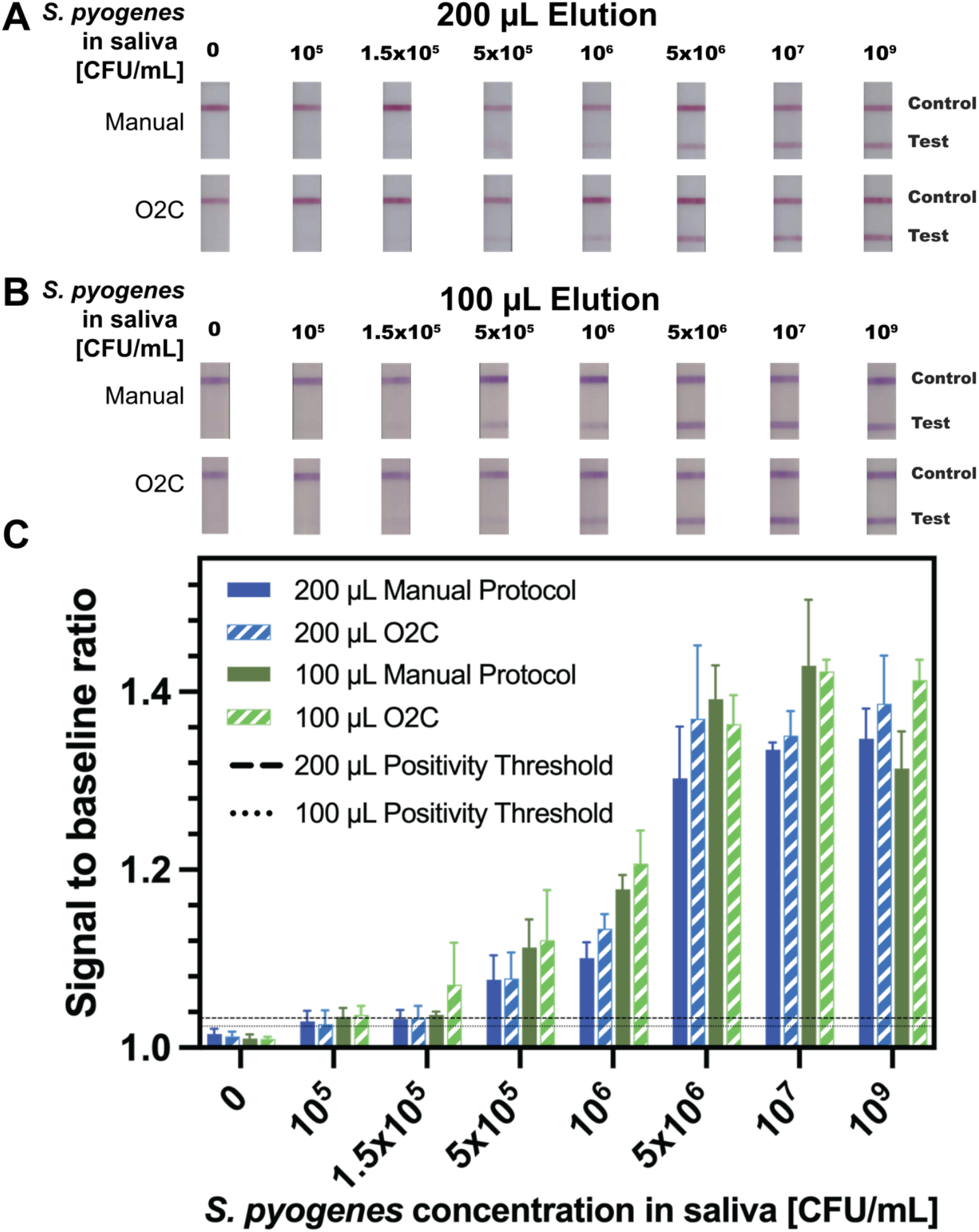
The microfluidic O2C procedure provides similar performance to the manual elution protocol. **A)** Lateral flow strip images were taken from GAS concentrations ranging from 0 CFU/mL to 10^9^ CFU/mL, with the manufacturer-specified limit of detection at 1.5×10^5^ CFU/mL. The manual protocol refers to the current RADT protocol^9^ with the CandyCollect substituted for the pharyngeal swab. O2C refers to the elution protocol performed with the microfluidic O2C platform. These results were obtained using the standard elution buffer volume of 200 µL. **B)** Lateral flow strips images taken from the same GAS concentrations as A with an elution buffer volume of 100 µL. **C)** Image analysis of the 200 µL and 100 µL dataset. Positivity threshold is the mean signal to baseline ratio + 3 standard deviations of the negative controls. Generally, the 100 µL elution with the O2C device produced the highest signal to baseline ratio at the different concentration levels with a signal to baseline ratio above the positivity threshold for all concentrations tested except 0.

Both elution methods also showed faint positive test lines at 1.5×10^5^ CFU/mL (the manufacturer-specified detection limit) with quantified test lines very close to the positive threshold.

To investigate the impact of sample pre-concentration on LFA results, we decreased the total elution volume to 100 µL. For both the manual and microfluidic methods, GAS concentrations between 5 ×10^5^ and 10^9^ CFU/mL again provided very clear test lines that were clearly above the positivity threshold **(Figure 3B and S2)**. At lower GAS concentrations (10^5^ – 10^6^ CFU/mL), test lines from 100 µL elution volume provided larger signals than the 200 µL elution, likely because of sample pre-concentration in the smaller volume. At higher GAS concentrations, the test line intensities were comparable across both elution volumes, likely because the signal had already saturated the test lines.

Taken together, our results suggest that the CandyCollect O2C microfluidic approach for sample elution provides comparable results to the traditional manual approach and can detect clinically relevant concentrations of GAS spiked into saliva. The O2C has the potential to enable a more integrated sample-to-answer workflow, where the user simply places a CandyCollect into the device, pushes a button, and gets LFA results. Ongoing work is focused on directly interfacing the LFA strip onto the O2C device to minimize risk with potentially infectious agents, integrating a reagent storage system for elution buffers, and conducting a usability assessment among minimally trained users prior to a clinical validation study. Our microfluidic approach also has the potential for spatial multiplexing and testing multiple LFA strips for different respiratory infections (e.g. COVID-19, Flu, RSV, etc).

Additional versions of Figure 3 with the data reorganized for elution volume and with incorporated data from the kit-provided swabs are shown in **Figures S3 and S4**, respectively.

## Conclusion

We developed the CandyCollect O2C microfluidic system that uses a 3D-printed sandwich with closed channels to extract liquid from the open channel CandyCollect saliva sampling device to simplify downstream testing with RADT kits. Our prototype represents a general class of device that can integrate open and closed microchannels to leverage the capabilities of both. Our results with GAS spiked into saliva samples show that the O2C provides similar or better performance to the manual elution method, while also concatenating multiple elution steps, automating mixing, and enabling easy manipulation of smaller sample volumes. The initial prototype shows excellent promise in being able to integrate user-friendly sampling with a reliable, rapid diagnostic platform. Further work will streamline device assembly and manufacturability, incorporate stored reagents, and automate the delivery of liquid to the LFA. We will also develop new prototypes that enable multiplexed detection of several respiratory pathogens within the same sample. The CandyCollect O2C microfluidic diagnostic platform has the potential to provide a child-friendly sampling method for consistent, hassle-free access to decentralized testing for minimally trained users.

## Materials & Methods

### Fabrication of the CandyCollect O2C System

#### Fabrication of CandyCollect device

CandyCollect devices (**Figure S5**) were milled from 2 mm thick sheets of polystyrene using a computer numerical control (CNC) Datron neo milling machine (GoodFellow, Cat# 235-756-86). Subsequently, the CandyCollect devices were sonicated with isopropanol (IPA) and 70% v/v ethanol (Fisher Scientific, Decon™ Labs, 07-678-004).^6,7^ Modified CandyCollect devices were created for the O2C system which had a 2 mm through-hole cut into the center of the spiral and a 0.75 mm inlet located on the right-hand side of the lollipop face.

#### Fabrication of the O2C device

The layers of the microfluidic device (**Figure S6**) were designed in Solidworks (Waltham, Massachusetts, USA) and 3D-printed with clear resin (Clear Microfluidics Resin V7.0a, CADworks3D, Toronto, ON, Canada) using a digital light processing printer (CADworks ProFluidics 285D, CADworks3D, Toronto, ON, Canada) which leverages a native 40 µm XY pixel resolution and an adaptive 28.5 µm XY pixel resolution. After printing, the devices were washed in an isopropanol (Sigma Aldrich, Cat#190764) bath, dried using a compressed nitrogen gun, and post-cured in a Formcure (Formlabs, Somerville, Massachusetts, USA) for 30 seconds on each side of the device.

The device ceiling layer (**Figure S6**) was trimmed from a 0.3 mm thick sheet of silicone rubber (Laimeisi, Shenzhen, China) using a commercial craft cutter machine (Silhouette Cameo 4).^17^ A 0.8 mm circular through-hole was cut into the silicone to align with the inlet of the CandyCollect device.

A finger pump was retrofitted from an e-cigarette cap^14^ (ELF bar 600 disposable Vape Pod). A two mm hole was punched through the top of the cap to allow air to escape. After surface treating the top layer, the reagent channel was covered with hydrophobic tape (ARseal™ 90697, Adhesives Research). A one mm outlet was left open at the end of the channel for the liquids to traverse into the CandyCollect microchannel.

#### Surface activation of CandyCollect device and reagent channel

The CandyCollect devices and reagent channel on the top layer of the device were activated by plasma treatment with oxygen using a Zepto LC PC Plasma Treater (Diener Electronic GmbH Plasma Treater, EbHausen, Germany). The following protocol is consistent with previous publications from our lab.^6,7^ In short, the chamber of the plasma treater will release enclosed atmospheric gas, in order to achieve a new pressure of about 0.20 mbar. Afterwards, the chamber is then surged with oxygen gas to establish a pressure of 0.25 mbar. Lastly, the chamber is then exposed to 70 W of voltage for a duration of five minutes.

### S. pyogenes culture

#### Liquid media preparation of Todd-Hewitt yeast (THY) broth & S. pyogenes maintenance

The *Streptococcus pyogenes* strain SF 370 (American Type Culture Collection, ATCC^®^, Cat#700294) was used for all experiments. Todd-Hewitt Broth (TH Broth) (BD Bacto™ TH broth, Fisher Scientific, Cat#DF0492-17-6), containing 0.2% yeast extract (United States Biological Corporation, Fisher Scientific, Cat# NC9796728) (THY) was used to culture *S. pyogenes*. Freeze-dried *S. pyogenes* was transferred from a frozen stock kept in a −80 C freezer into a 15 mL conical tube (Fisher Scientific, Falcon™, Cat# 14-959-49B) containing 10 mL of THY media. The culture was then incubated overnight at 37 ^°^C supplied with 5% CO2.^9^

#### S. pyogenes agar maintenance

*S. pyogenes* was maintained by streaking bacteria onto the agar plates with a sterile disposable inoculating loops (Globe Scientific, Fisher Scientific, Cat# 22-170-201). The agar plates were incubated overnight at 37 ^°^C supplied with 5% CO2. Agar plates were stored at room temperature for up to 7 days. An individual colony from the agar plate was used to inoculate with 10 mL of THY liquid media confined in a 15 mL conical tube (Fisher Scientific, Falcon™, Cat# 14-959-49B) and cultured overnight as above.^6,7^

### Capture, elution, and detection of *S. pyogenes* in manual vs microfluidic procedure

#### Suspension of S.pyogenes in filtered pooled human saliva

To estimate the concentration of S. pyogenes in liquid culture media, optical density at 600 nm (OD600) was measured using a Visible 721-Vis Spectrometer (Vinmax). To achieve a pellet of *S. pyogenes*, the liquid culture was centrifuged for 5 minutes at 10,000 rpm. Afterwards the *S. pyogenes* pellet was resuspended with filtered pooled human saliva (Innovative Research, Cat# IRHUSL50ML; filtered using a 0.22 μm filter). Finally, serial dilutions were performed to achieve the desired *S. pyogenes* concentration.^9^

#### Capture of S. pyogenes on the open fluidic channels on the CandyCollect device

50 μL of *S. pyogenes* in saliva were incubated on the CandyCollect device for 10 mins with the following concentrations: 1.0×10^7^, 5.0×10^6^, 1.0×10^6^, 5.0×10^5^, 1.5×10^5,^ and 1.0×10^5^ colony forming units (CFU)/mL; with the addition of a positive control, *S. pyogenes* at 1×10^9^ CFU/mL suspended in liquid media, and a negative control, filtered pooled saliva.^9^

#### Extraction of surface antigen from S. pyogenes through manual procedure

After the 10 min incubation time, the CandyCollect device was transferred into a 15 mL test tube containing the elution reagents: 2.0 M Sodium Nitrite (Reagent A) and 0.4 M Acetic Acid (Reagent B). Previous studies reported that these reagents produced the highest test line intensity compared to other reagents.^9^ Afterwards, the test tube was shaken for 15 s to ensure that the liquid had adequate contact with the CandyCollect microchannel and left to sit for 45 s.

#### Extraction of surface antigen from S. pyogenes using the CandyCollect O2C system

The CandyCollect device was placed into the inset of the bottom layer. The device ceiling layer was placed on top of the CandyCollect with the inlet on the CandyCollect and the through-hole in the ceiling layer aligned. The top layer was positioned on the bottom layer and the device was clamped tightly together with binder clips. The entire assembly took around one minute, matching the timing of the manual procedure. The finger pump was placed in the inset of the top layer and the liquid was eluted into a waiting petri dish (Fisher Scientific, Corning, Cat#07-200-83) upon a single press.

### Lateral flow immunoassay strip readout and data analysis

#### Examination by eye of lateral flow immunoassay strip

A lateral flow immunoassay (LFA) strip from the Areta kit was saturated with the eluate from either the manual elution procedure or the O2C. Following manufacturer’s instructions^18^ to ensure enough eluted solution was added to the LFA, we held the strip in the eluted solution until wicking was observed at the start of the nitrocellulose membrane and then removed and set aside in a petri dish (Fisher Scientific, Cat# 165218) with a timer set for 18 min. After 18 min, we examined the strips in lab by eye and recorded the results which are valid from 10 min to 20 min after the test strips have been removed from the elution solution. Lastly, strips were scanned (Hewlett-Packard, HP OfficeJet Pro6978, SN# TH0AK4N0YT) at 600 dpi and darkness = 7 to a JPEG format (**Figure S7**).^9^

#### Quantification of signal on Areta lateral flow immunoassay strip via optical analysis

We used a custom image analysis Python script to analyze the scanned LFA strips, using an algorithm published in previous work. The scanned images were cropped to 25 × 100-pixel images from the original full-color images obtained. The images were converted to monochrome, inverted, and profiles of digital numbers against the pixel location were obtained by averaging the rows for each strip. The code denotes the peak signal values from the test line against location on the lateral flow strip. To establish baseline signal, two regions-of-interest (ROIs) spanning between 5 to 10 pixels were manually selected, approximately 30 pixels from both sides of the peak. The average digital number from these two ROIs is calculated to compute the baseline. To obtain the signal-to-baseline ratio (SBR), the ratio of the peak signal of the test line from the profile is divided by the baseline.

The positivity threshold is a value found from the SBR, above which the test results are deemed positive. It is derived from image analysis of the three negative controls collected from each individual experiment and is calculated as such:

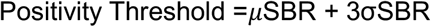

where μSBR is the average signal-to-baseline and σSBR is the standard deviation signal-to-baseline of the three negative controls. The python script finds the negative signal by analyzing a segment of the profile located a specific distance from the negative control line peak, a distance which is informed by the test line peak locations in positive test strips. This approach ensures that the location of the segment accurately reflects the expected position of the negative control test line.

See SI for expansion of materials and methods.

## Acknowledgments

We want to acknowledge the M.J. Murdock Diagnostic Foundry for Translational Research at UW for allowing us to use equipment. We would also like to thank Dr. Lucas Meza and the Meza Lab for access to instruments. We are grateful for funding from the National Institute of Health (NIH) R35 grant #GM128648. The content is solely the responsibility of the authors and does not necessarily represent the official views of the National Institutes of Health or other funding sources.

## Author Contributions

K.M.L., A.B.T, E.B., and A.O.O. conceptualized the initial design integrating the CandyCollect with a microfluidic device. K.M.L., J.C.S., A.B.T., and A.O.O. conceptualized the wet lab studies to test the platform with spiked saliva. J.C.S. milled the CandyCollect devices. K.M.L. 3D printed and craft cut the microfluidic device and ceiling device. K.M.L., R.R.A., J.A.T., and C.C. designed the microfluidic device in CAD. K.M.L. and J.A.T. performed the initial device proof-of-concept tests. J.C.S. and K.M.L. designed the biological experiments, conducted the biological experiments, and performed data collection. A.B.T. and A.O.O. advised on study design and execution. J.C.S., T.R.R., M.M.C., A.O.O., E.B., and A.B.T. developed and refined data analysis methods for quantifying RADT results. All authors analyzed and interpreted data. K.M.L. and A.O.O. wrote sections of the manuscript. K.M.L. created all visual work presented in the paper. All authors revised and approved the manuscript.

