## Supplementary for "CandyCollect Open to Closed (O2C) Microfluidic System for Rapid and User-Centric Detection of Group A Streptococcus"

**Link to repository with CAD files of O2C system and ceiling device:**
https://doi.org/10.5281/zenodo.14681284
**GitHub with image analysis code:**
https://github.com/timrobinson/UW-LFA-Analysis.git
**Figure S1:** Images of all lateral flow strips from the 200 µL elution experiment
**Figure S2:** Images of all lateral flow strips from the 100 µL elution experiment
**Figure S3:** Graphs reorganized by elution volume
**Figure S4:** Graph of kit-provided swab data incorporated into 200 µL and 100 µL elution data
**Figure S5:** Engineering drawing of CandyCollect used with O2C system
**Figure S6:** Engineering drawing of CandyCollect O2C system
**Figure S7:** Scanner optimization
**Extended Materials and Methods Section**

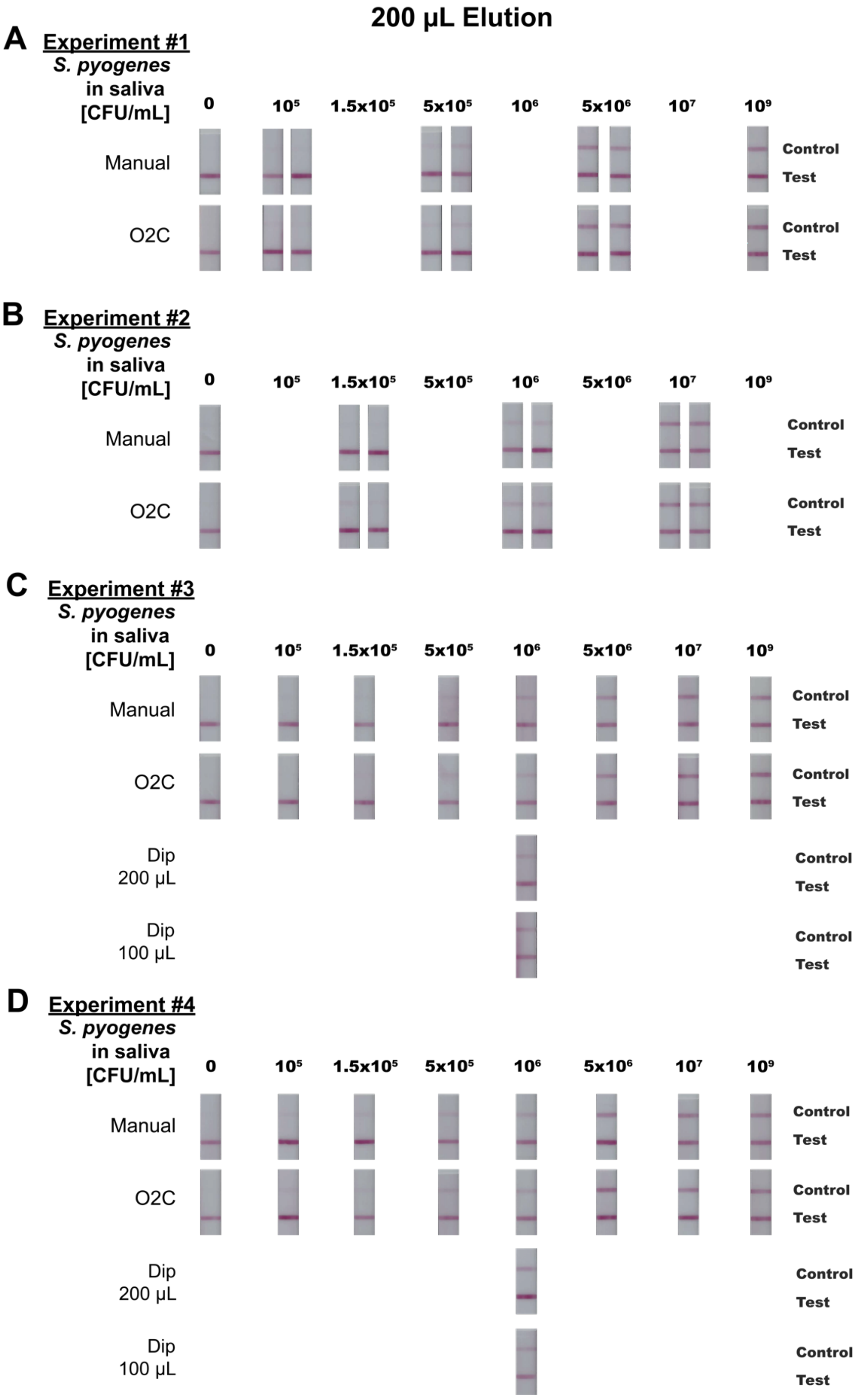


**Figure S1:** Images of lateral flow strips from four individual 200 µL elution experiments to compare the manual and O2C procedures. These images were used to obtain the image analysis results depicted in figure 3. The images support our findings that the manual and O2C procedures produced comparable results across all concentrations.

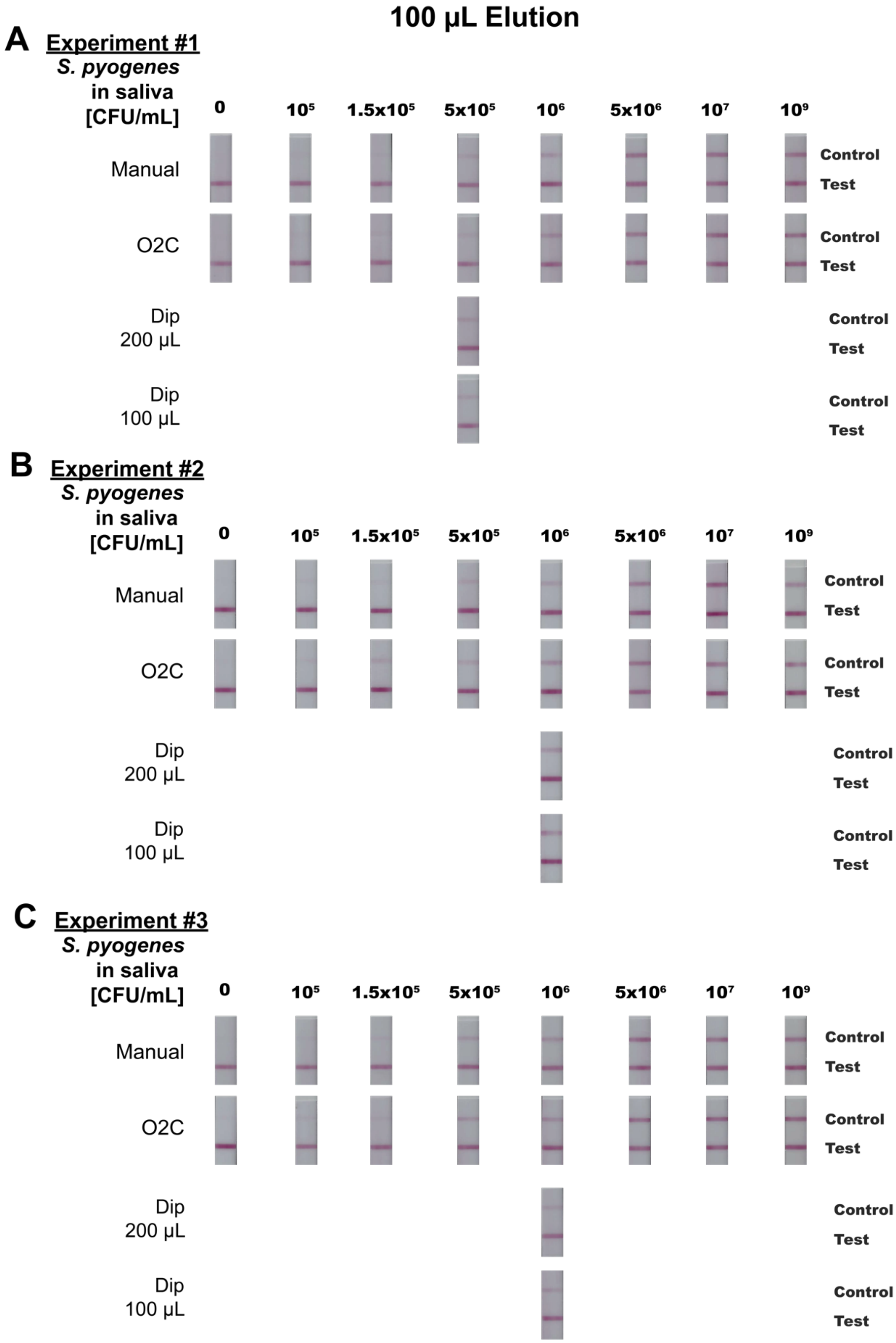

**Figure S2:** Images of lateral flow strips from three individual 100 µL elution experiments to compare the manual and O2C procedures. These images were used to obtain the image analysis results depicted in figure 3. The images support our findings that the manual and O2C procedures produced comparable results across all concentrations. We also see more prominent test lines at lower concentrations in the 100 µL lateral flow strips compared to the 200 µL lateral flow strips highlighting that effects of using a lower elution volume.

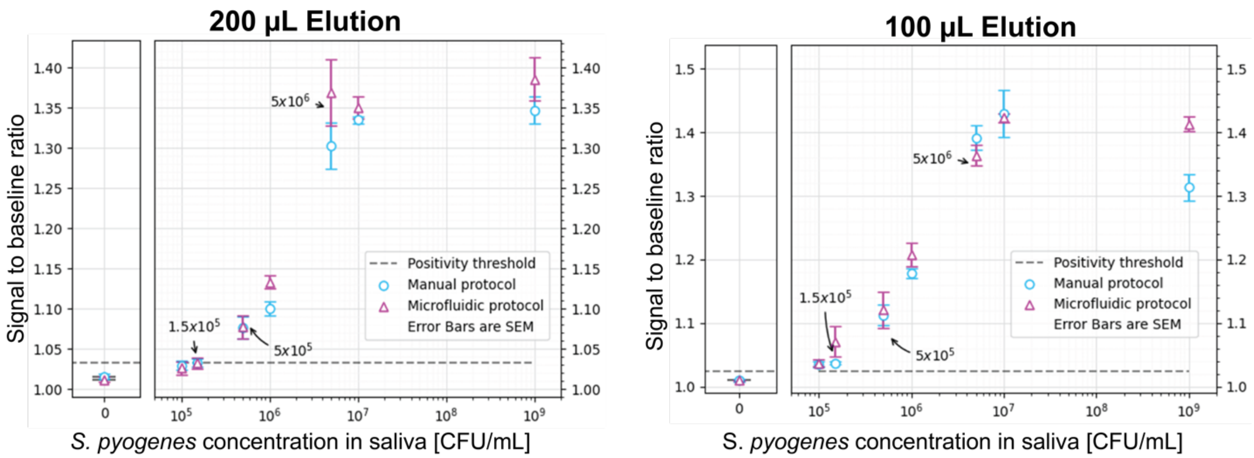


**Figure S3:** Graphs of 200 µL elution image analysis data and 100 µL elution image analysis data in individual plots. This is the same data in figure 3 but reorganized into individual plots to demonstrate the differences between manual and microfluidic protocol for each individual elution.

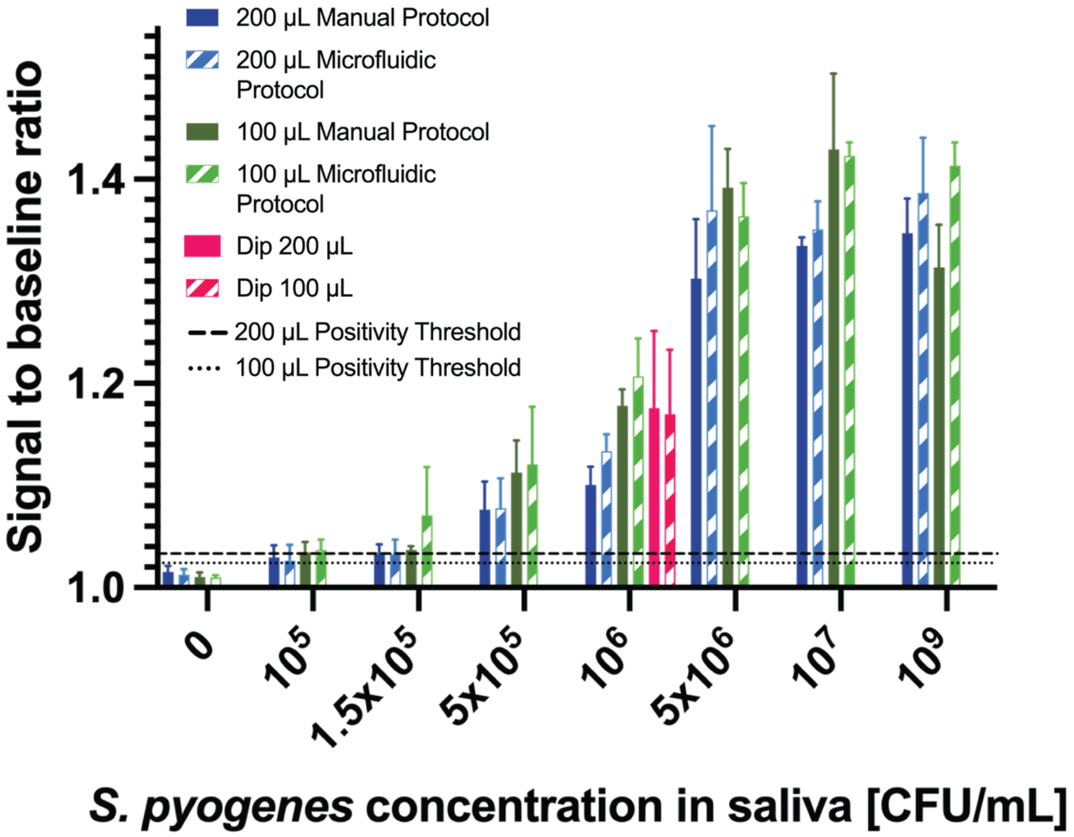


**Figure S4:** Graph of dip data integrated into current elution data. 'Dip' refers to the method of using the kit-provided swab and dipping the swab into the saliva sample prior to performing the elution steps. This was used a gold standard method when previously evaluating the manual method.[1] In this paper, the goal is to compare the manual and microfluidic method, so the inclusion of the dip data here is for completeness. The dip method was tested at both 200 µL and 100 µL elution volumes and produced comparable results to the manual and microfluidic methods.

### Engineering Drawings

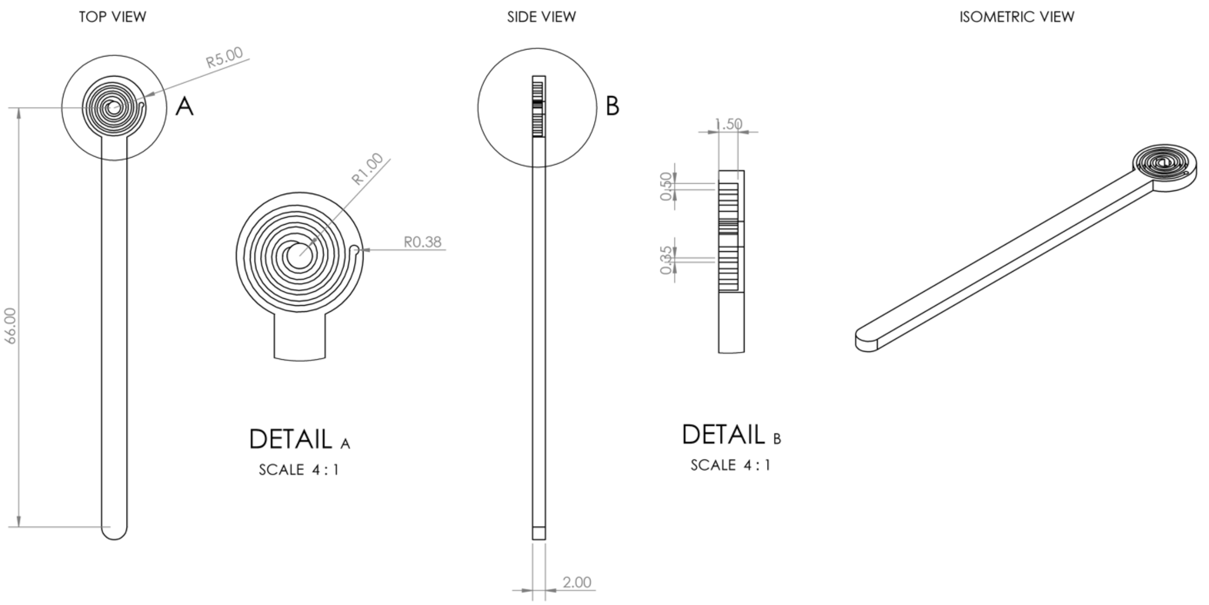


**Figure S5:** Engineering drawing of CandyCollect device used with O2C system in the experiments (Figure 3). The device has a hole in the center to enable elution of the liquid. All dimensions are in mm.

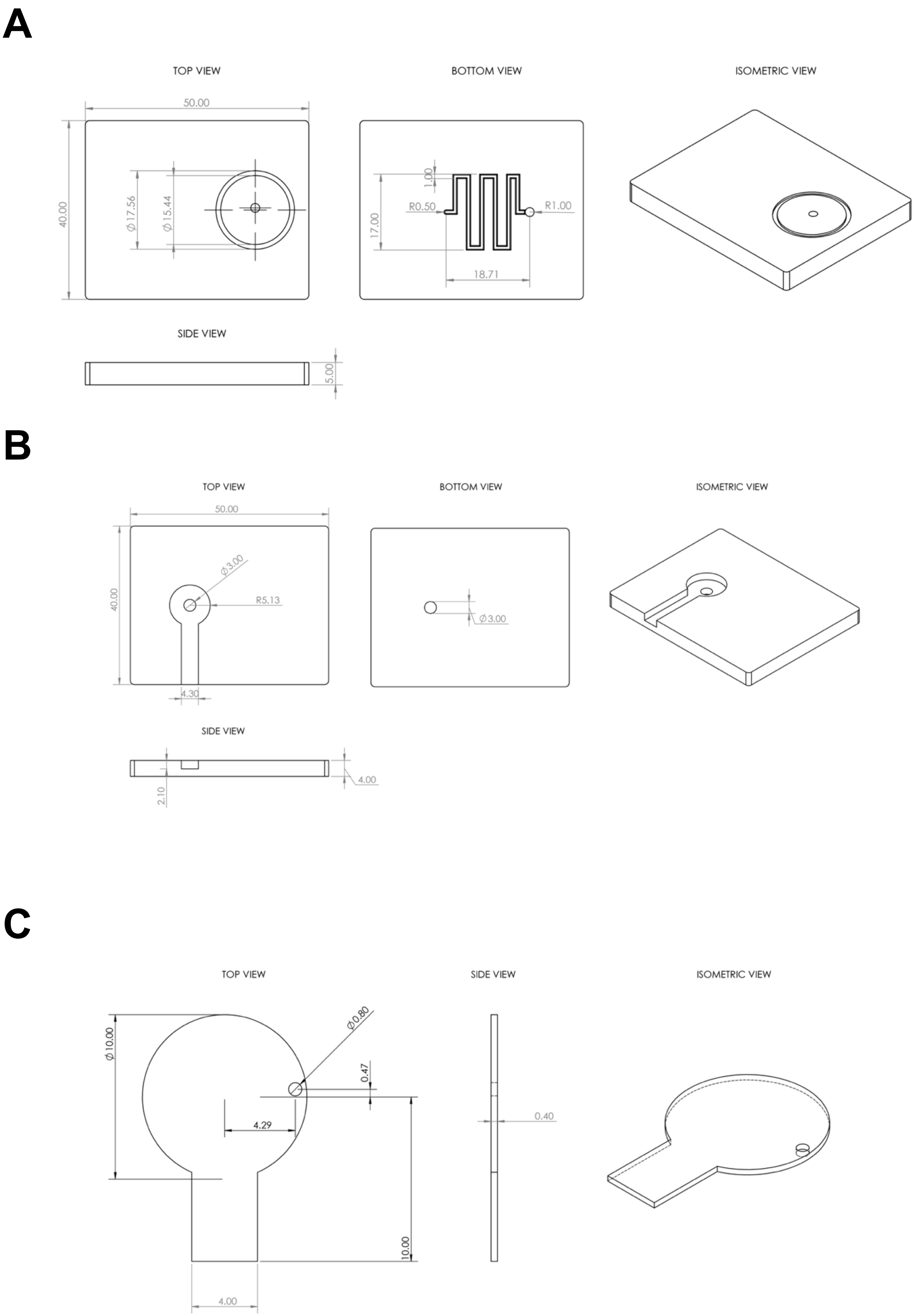

A
TOP VIEW
BOTTOM VIEW
ISOMETRIC VIEW
50.00
40.00
Ø17.56
Ø15.44
1.00
R0.50
R1.00
17.00
18.71
SIDE VIEW
5.00
B
TOP VIEW
BOTTOM VIEW
ISOMETRIC VIEW
50.00
Ø3.00
R5.13
40.00
4.30
Ø3.00
SIDE VIEW
4.00
2.10
C
TOP VIEW
SIDE VIEW
ISOMETRIC VIEW
Ø10.00
Ø0.80
0.47
4.29
0.40
10.00
4.00

**Figure S6:** Engineering drawing of CandyCollect O2C system used in experiments (Figure 3). A) The top layer of the O2C system contains the microfluidic channel and inset for the finger pump. B) The bottom layer of the O2C system has an inset for the CandyCollect and contains a hole for liquid to be eluted into a reservoir. C) The device ceiling layer which has a hole for liquid to move into from the top microchannel. All dimensions are in mm.

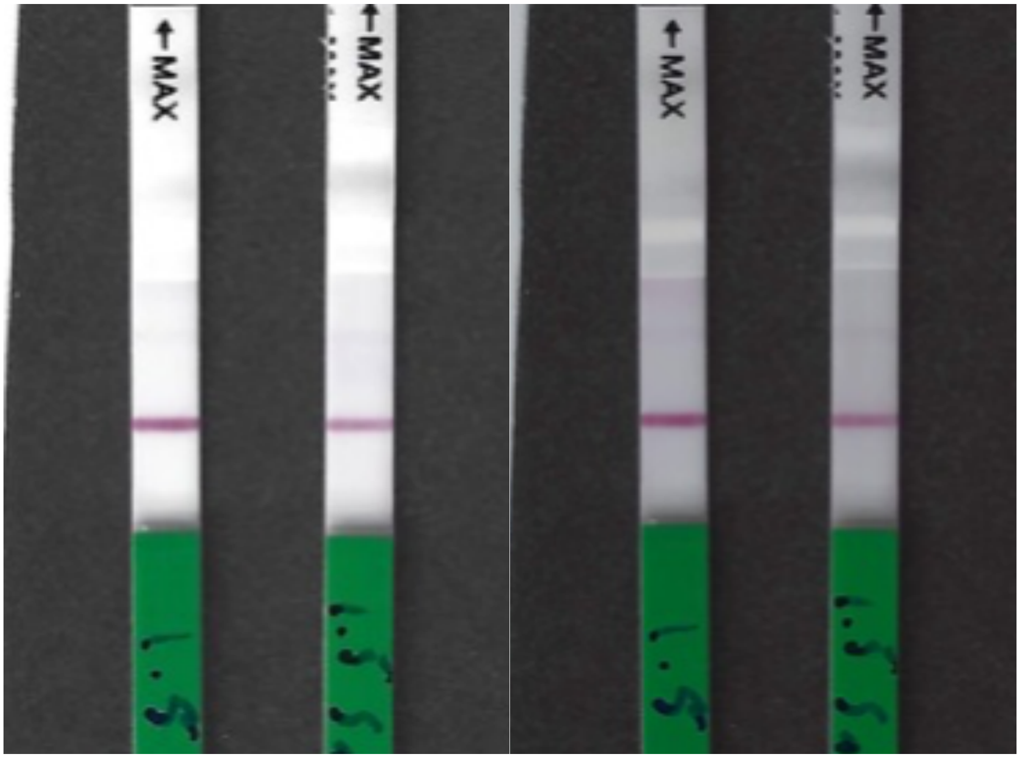


**Figure S7:** Optimization of the scanner for lateral flow strip. We optimized the commercial scanner to obtain high-quality photos of our lateral flow strips. The default settings of the scanner are shown on the left with a DPI of 300 and a darkness setting of 5. We optimized the settings to obtain more contrast between the test line, thereby resulting in an image that was more representative of what we were seeing by eye. The photo on the right represents the optimized settings with a DPI of 600 and a darkness of 7.

**Extended Materials and Methods Section**

*Preparation of* S. pyogenes *in filtered pooled human saliva (Reproduced from Sanchez et al.*[1]*)*

Optical density at 600 nm (OD600) was measured to estimate the concentration of *S. pyogenes* growing in liquid culture media using a Visible 721-Vis Spectrophotometer (Vinmax). Liquid culture was centrifuged to pellet cells for 5 min at 10000 rpm. The *S. pyogenes* pellet was resuspended in filtered pooled human saliva (Innovative Research, Cat# IRHUSL50ML; filtered using a 0.22 μm filter. To achieve desired concentrations of *S. pyogenes*, serial dilutions were performed. See the section below for OD600 to CFU/mL conversion.

*Converting between OD600 absorbance measurements of S. pyogenes to CFU/mL (Reproduced from Sanchez et al.*[1]*)*

To generate OD600 to CFU/mL conversion factor, first, the OD600 of a bacterial culture was measured. Second, a small volume (50-100 μL) of serial dilution of bacterial culture was spread plated on agar plates. After overnight culture, viable colonies on each agar plate were counted. Combined with plating volume and dilution factor, CFU/mL was calculated. Then OD600 measurement was correlated with CFU/mL measurement. The same correlation/conversion factor was used in all experiments.

For further instruction on the image analysis process please refer to the supplementary of Sanchez et al.[1]
